# Ethanol-Mediated Compaction and Crosslinking Enhance Mechanical Properties and Degradation Resistance While Maintaining Cytocompatibility of a Nucleus Pulposus Scaffold

**DOI:** 10.1101/333179

**Authors:** Joshua D. Walters, Sanjitpal S. Gill, Jeremy J. Mercuri

**Author notes:** **Corresponding Author:** Jeremy Mercuri, Ph.D., 313 Rhodes Engineering Research Center, Clemson, SC, 29634, (email). (Joshua D. Walters), (Sanjitpal S. Gill), (Jeremy J. Mercuri).

## Abstract

Intervertebral disc degeneration is a complex, cell-mediated process originating in the nucleus pulposus (NP) and is associated with extracellular matrix catabolism leading to disc height loss and impaired spine kinematics. Previously, we developed an acellular bovine NP (ABNP) for NP replacement that emulated human NP matrix composition and supported cell seeding; however, its mechanical properties were lower than those reported for human NP. To address this, we investigated ethanol-mediated compaction and crosslinking to enhance the ABNP’s dynamic mechanical properties and degradation resistance while maintaining its cytocompatibility. First, volumetric and mechanical effects of compaction only were confirmed by evaluating scaffolds after various immersion times in buffered 28% ethanol. It was found that compaction reached equilibrium at ∼30% compaction after 45 min, and dynamic mechanical properties significantly increased 2-6x after 120 min of submersion. This was incorporated into a crosslinking treatment, through which scaffolds were subjected to 120 min pre-compaction in buffered 28% ethanol prior to carbodiimide crosslinking. Their dynamic mechanical properties were evaluated before and after accelerated degradation by ADAMTS-5 or MMP-13. Cytocompatibility was determined by seeding stem cells onto scaffolds and evaluating viability through metabolic activity and fluorescent staining. Compacted and crosslinked scaffolds showed significant increases in DMA properties without detrimentally altering their cytocompatibility, and these mechanical gains were maintained following enzymatic exposure.

## Introduction

Low back pain (LBP) is the leading cause of Years Lived with Disability in all developed countries,^1^with a lifetime prevalence of 70-84%.^2^ LBP has severe economic ramifications on the individual and societal levels, leading to premature workforce departure, short/long-term fiscal losses,^3^ and cumulative yearly costs exceeding $100 billion in the US.^4^ The most common diagnosis for patients experiencing LBP is intervertebral disc (IVD) degeneration (IDD).^5^

IVDs are avascular, fibrocartilaginous structures connecting adjacent vertebrae in the spinal column. Each IVD comprises an aggrecan-rich core known as the nucleus pulposus (NP), 15-25 concentric sheets of aligned collagen type I encircling the NP known as the annulus fibrosus (AF), and thin layers of hyaline cartilage covering the cranial and caudal surfaces of the IVD known as the cartilage endplates (CEP).^6^ Adjacent vertebrae interface with the IVD through the CEP, which sequesters the NP within the disc while allowing for nutrient exchange with vertebral blood vessels.^7^ The encapsulation of the NP and its anionic, dense matrix impart triphasic time-dependence to its mechanical properties, allowing it to dissipate loads via fluid exudation and prevent harmful stress concentrations in AF and CEP through hydraulic pressurization.^6,8,9^

IDD is a multi-factorial, cell-mediated process originating in the NP in which normal tissue turnover becomes imbalanced, favoring degradation over formation.^10–12^ This degradative environment is known to include metalloproteinases (MMPs) and a disintegrin and metalloproteinase with thrombospondin motifs (ADAMTS) – catabolic enzymes that respectively degrade the two main constituents of the IVD, collagen and aggrecan.^13–15^ This disparity between anabolism and catabolism results in altered extracellular matrix (ECM) composition, loss of structural morphology, and diminished spinal function.^16^ Treatments for IVD disorders range from palliative therapies, such as physical therapy and pharmacological pain management,^17,18^ to surgical interventions, such as intradiscal electrothermal therapy, spinal fusion, or IVD arthroplasty.^19,20^ However, the efficacy of current treatments remains controversial, particularly in regards to alleviating chronic discogenic pain and maintaining biomechanical function.^21–24^ There is a need for novel therapies capable of restoring IVD function and maintaining native tissue structure while providing long-term regeneration. Incorporating stem cells into a mechanically robust, tissue-engineered scaffold could satisfy this need.

Decellularization is a provocative tissue engineering strategy due to its ability to remove immunogenic components from tissue while retaining native ECM composition and organization.^25^ Past work has shown that cartilage-mimetic scaffolds formed through decellularization can induce stem cells to adopt NP-like cell phenotypes,^26^and that ECM composition and density of the NP gives rise to its unique poroviscoelastic mechanical properties.^27–29^ These suggest that decellularization could produce NP replacement scaffolds with compositions similar to the native ECM, imbuing them with the intrinsic ability to induce NP-like cell phenotypes while possessing comparable mechanical properties. Decellularized NP therapies have been previously reported, but many are implemented as injectable derivatives of acellular ECM, ^30–32^ and have not yet been evaluated for mechanical function post-injection. Decellularization of intact human NP was reported by Huang et al, who demonstrated devitalization of the tissue through the absence of nuclei within lacunae and the preservation of native ECM components.^33^ However, quantification of DNA content showed no difference between native and decellularized tissues, which could induce an adverse host reaction in response to residual exogenous DNA,^34,35^ and the decellularized tissue was not mechanically characterized. Mercuri et al successfully demonstrated the decellularization of porcine NP and its ability to support stem cell differentiation towards an NP-like phenotype in the absence of exogenous growth factors, however the mechanical properties were within the lower margin of human NP values.^36,37^

Bovine caudal NPs have shown biochemical and microarchitectural similarities to human NP.^28,37^ We have previously described an acellular bovine NP (ABNP) which possessed biomimetic ECM composition and supported cell seeding,^38^ and have recently shown its ability to partially restore spine kinematics using a bovine explant functional spinal unit model when used in tandem with an annulus fibrosus repair patch.^39^ There remains a need to optimize the ABNP prior to its implementation as a therapeutic for IDD to ensure it is mechanically competent when confronted with enzymatic degradation.

A ubiquitous method of fabricating and fortifying biomaterials for implantation is using 1– ethyl–3-(3-dimethylaminopropyl) carbodiimide hydrochloride and N-hydroxysuccinimide (EDC/NHS) crosslinking to control degradation kinetics and mechanical properties.^40^ EDC/NHS is a zero-length crosslinking treatment conjugating free amines and carboxyl groups,^41^ and it has received attention because it is not incorporated into the scaffold during the crosslinking reaction, allowing cytotoxic crosslinkers and byproducts to be rinsed from the scaffold prior to implantation.^41^ Additionally, using ethanol (EtOH) as a co-solvent has been shown to enhance EDC/NHS crosslinking, optimally at 28% EtOH,^42^and ethanol is known to shrink tissues during histological processing.^43^ We hypothesized that adding EtOH as a co-solvent would compact the ABNP during crosslinking, independently contributing to enhanced scaffold properties. Thus, the objectives of this research were to incorporate ethanol as a co-solvent for crosslinking the ABNP and to evaluate the effects of compaction and crosslinking on the dynamic mechanical properties, degradation resistance, and cytocompatibility.

## Materials and Methods

### Decellularization Procedure

Bovine coccygeal discs (CC1/2-CC3/4) were decellularized to create acellular bovine NP (ABNP) as described previously.^38^ Briefly, fascia was excised from bovine tails (Publix, Lakeland, FL) and then IVDs were dissected from adjacent vertebrae. The NP was isolated using an 8mm biopsy punch and decellularized using a series of detergents, ultrasonication, and nucleases. In preparation for mechanical testing, scaffolds were snap-frozen, trimmed into cylinders (Ø4 mm; mean height: 3.26 ± 0.13 mm), and thawed in saline. Scaffolds used for cell culture were sterilized in 0.1% Peracetic Acid (Sigma-Aldrich, St. Louis, MO) in PBS for 2 hr, and then neutralized overnight in Dulbecco’s Modified Eagle Medium (DMEM: Fisher Scientific, Waltham, MA) containing 50% fetal bovine serum (FBS: Atlanta Biologicals, Flowery Branch, GA) and 1% antibiotic/-mycotic (AB/AM: Fisher Scientific).

### Volumetric Confirmation of Ethanol-Mediated ABNP Compaction

A preliminary study was performed to confirm scaffold compaction after immersion in 28% ethanol buffered with 50 mM 2-(N-Morpholino) ethanesulfonic acid (MES: Merck Millipore, Burlington, MA). Scaffolds (n=9) were imaged from the top and side prior to immersion, and post-immersion at 15, 30, 45, 60, 90, 120, and 180 min. Using ImageJ (National Institutes of Health, Bethesda, MD), the height and diameter of each scaffold was measured at each time point, and changes were calculated as a percentage of the initial volume.

### Dynamic Mechanical Analysis of Ethanol-Mediated ABNP Compaction

To isolate the mechanical effects of ethanol-mediated compaction, a preliminary unconfined dynamic mechanical analysis (DMA) study was performed on a Bose Electroforce 3200 test frame (TA Instruments, New Castle, DE) equipped with a saline bath at 27°C. ABNP (n=12) were equilibrated at 4% compressive strain for 500 sec followed by 15 cycles from 0% to 8% compressive strain at 1 Hz. After re-hydrating for 120 minutes in saline, samples were immersed in 28% EtOH buffered with MES for 30, 60, or 120 min (n=3/timepoint), and then subjected to an identical loading regimen. Stiffness, storage modulus and loss modulus were quantified for all DMA testing using Wintest7 DMA Analysis Software (TA Instruments).

### ABNP Crosslinking

Since preliminary testing showed that MES-buffered 28% ethanol induced significant changes in scaffold volume and all DMA properties after immersion for 120 min, this was incorporated into a crosslinking treatment whereby samples underwent 120 min of compaction prior to crosslinking. (Figure 1) Crosslinking was performed for 4 hr using 30 mM EDC (Fisher Scientific) and 6 mM NHS (Alfa Aesar, Haverhill, MA) in MES buffer at 5.5 pH in three different conditions: no ethanol co-solvent (MES Only), 28% ethanol co-solvent (EtOH/MES), or 120 min pre-compaction in buffered 28% ethanol co-solvent followed by the addition of EDC/NHS (Comp+EtOH/MES). Native bovine NP (bNP) and unaltered ABNP were used as controls.

**Figure 1.**
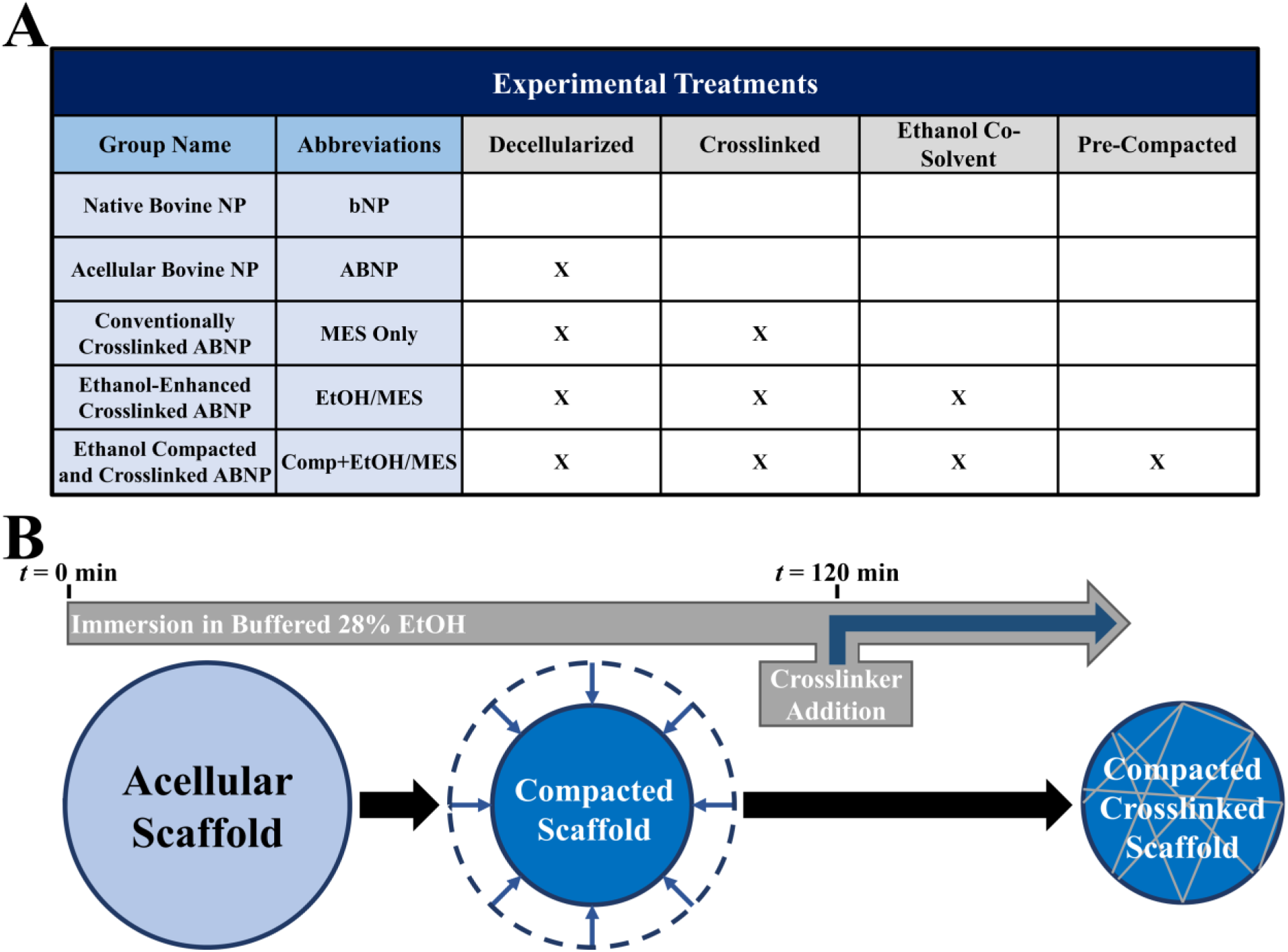
Experimental Design. **A**) Table depicting experimental study group name, abbreviated identifiers and descriptors. **B**) Schematic overview illustrating the proposed ethanol-mediated compaction and crosslinking approach for NP scaffolds.

### Compressive Dynamic Mechanical Analysis of Crosslinked and Compacted ABNP

To evaluate the dynamic mechanical changes after ethanol-mediated compaction and crosslinking, scaffolds (n=6/group) underwent DMA using a Bose Electroforce 3200 equipped with a saline bath at 27°C. Unconfined samples were equilibrated at 4% compressive strain for 500 sec, followed by 15 cycles from 0% to 8% compressive strain at 0.01 Hz, 0.1 Hz, 1 Hz, and 10 Hz, with a 500 sec equilibration at 4% prior to each waveform.

#### Compacted and Crosslinked ABNP Resistance to Accelerated Enzymatic Degradation

Following DMA, samples were lyophilized for enzymatic degradation studies using ADAMTS-5 and MMP-13, which have been shown to be important mediators of IVD degeneration. To simulate an accelerated degenerative environment, 50x high concentrations than those quantified by Mern et al were used.^44^ Samples (n=3/enzyme/group) were exposed to 1xTris-buffered saline containing 0.05% Brij-35, 5 mM CaCl_2_ and either recombinant human ADAMTS-5 (Fisher Scientific) at 0.20485 µg/mL or recombinant human MMP-13 (Fisher Scientific) at 0.02215 µg/mL. After 30 days of incubation at 37°C with daily agitation, scaffolds were re-lyophilized and their dry-weights were obtained to determine the percent mass loss. Scaffolds were then re-hydrated for 48 hr in PBS and then re-subjected to the DMA regimen.

### Cytocompatibility Testing of Crosslinked and Compacted ABNP

Each scaffold (Ø6 mm × 2-4 mm height; n = 4/group for ABNP, MES Only, EtOH/MES, Comp+EtOH/MES) was seeded with 1.0×10^6^ (passage 6) human Adipose Derive Stem Cells (hADSCs: Fisher Scientific) in 150μL of media. To minimize injection tract size, cell suspension was administered via multiple 25µL injections through a syringe tipped with a 29G needle, which we have previously shown to have no detrimental effects on cell viability.^38^ To allow for attachment, scaffolds were incubated for 2 hr at 37°C and 5% CO_2_ prior to the addition of cell culture media consisting of DMEM supplemented with 10% FBS and 1% AB/AM. Cell metabolic activity was assessed using Alamar Blue (Fisher Scientific) fluorescence at t = 1, 3, and 7 days, normalized to day 1 values. To confirm cell viability at day 10, scaffolds were bisected either sagitally or transversely (n=1/group/imaging plane) for Live/Dead fluorescent staining (Biotium, Fremont, CA) according to manufacturer instructions. Fluorescent images were acquired using an EVOS Fluorescent Imaging System (AMF4300: Thermo Fisher Scientific, Waltham, MA).

### Statistics

Results were expressed as mean ± standard error of the mean (SEM), with statistical analysis performed using Prism 7 (Graphpad Software, La Jolla, CA) and significance (p≤0.05) denoted by “*”. Volumetric changes were compared using one-way Repeated Measures ANOVA with Tukey’s *post hoc* pairwise comparison. DMA parameters for the preliminary compaction testing were compared to pre-immersion values (t = 0 min) using *Fisher’s LSD.* Subsequently, all DMA comparisons were performed within respective frequencies to the unaltered ABNP using *Fisher’s LSD.* Student’s t-test was used to compared DMA results before and after enzymatic degradation, with significance (p≤0.05) denoted by “†”. Percent mass loss was analyzed using one-way ANOVA with Dunnet’s *post hoc* comparison to the ABNP. Alamar Blue Fluorescence was analyzed using one-way ANOVA with Dunnet’s comparison to the ABNP within respective days.

## Results

### Ethanol-Mediated Compaction of ABNP

To confirm the ability of ethanol to mediate compaction of the ABNP, changes in scaffold volume were determined following various immersion times in 28% ethanol (Figure 2). Volumetric equilibrium was defined as no significant difference with respect to any previous timepoint. Volume was significantly reduced to 78.4±2.4% after 15 min (p<0.0003; 0 vs. 15 min) of compaction and equilibrium was reached at 69.4±1.9% after 45 min (p<0.0212; 15 vs. 45 min), after which time no significant changes in volume were found. This degree of volumetric compaction was re-confirmed after incorporation into a crosslinking treatment (data not shown). In addition to the volumetric changes, immersion in buffered 28% ethanol induced changes in the dynamic mechanical properties of the ABNP. Scaffold stiffness significantly increased from 45.3±8.9 mN/mm to 105.0±30.1 mN/mm after 60 min (p<0.0126) of immersion and to 118.3±18.0 mN/mm after 120 min (p<0.0036) (Figure 3A). Significant increases were also observed in the storage and loss moduli (Figure 3B and 3C), which respectively increased after 120 min from 8.0±1.1 kPa to 40.5±14.7 kPa (p<0.0003) and 2.5±0.3 kPa to 14.0±4.9 kPa (p<0.0003).

**Figure 2.**
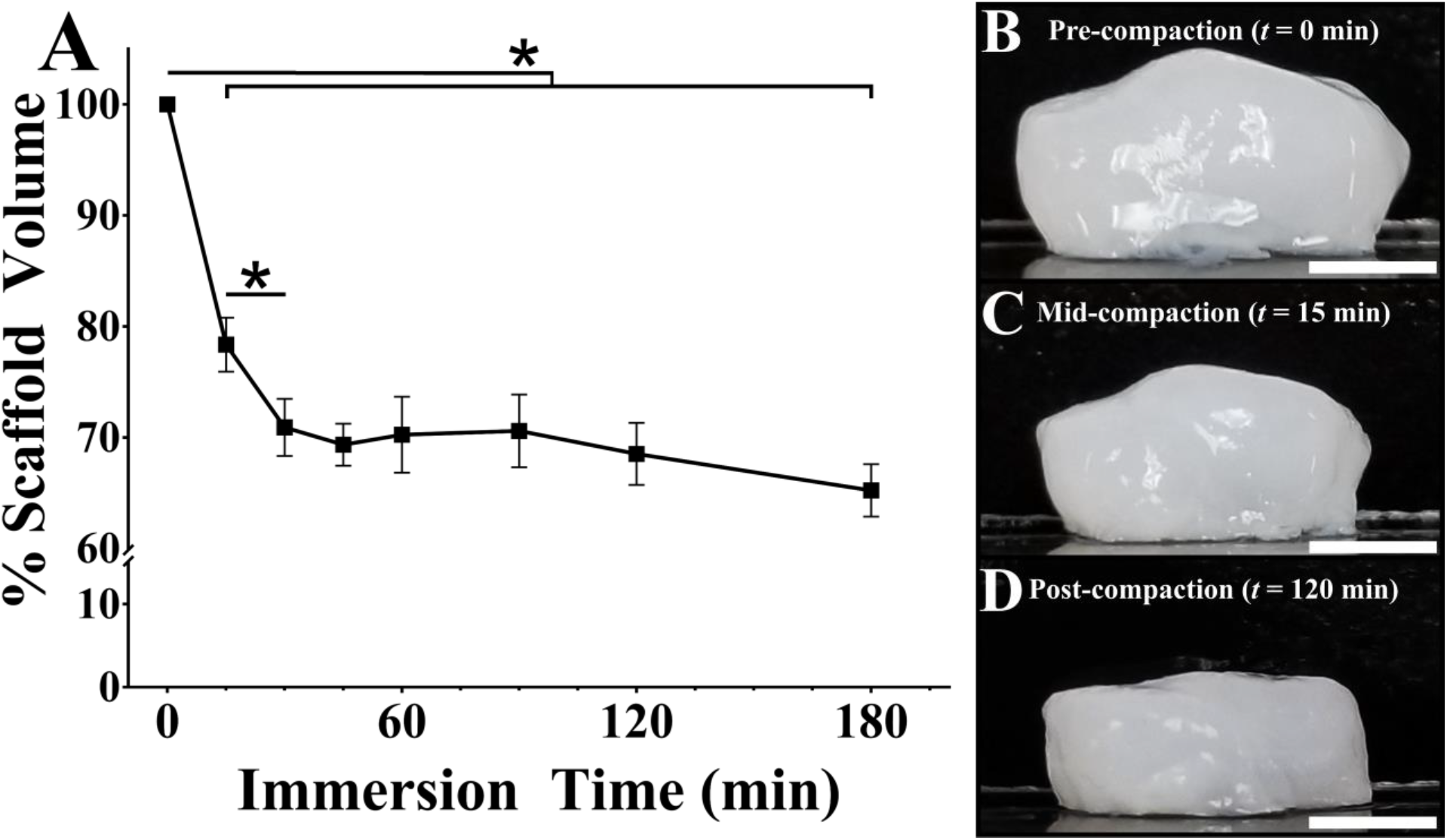
Effect of Ethanol Compaction on Scaffold Volume. **A**) Graph depicting the change in scaffold volume with respect to submersion time in 28% ethanol. Lines connecting study groups indicate statistical differences (p≤0.05). Representative macroscopic images of ABNP scaffold **B**) prior to (t = 0 minutes), **C**) during (t = 15 minutes) and **D**) after (t=120 minutes) of submersion in 28% ethanol (bar = 5 mm).

**Figure 3.**
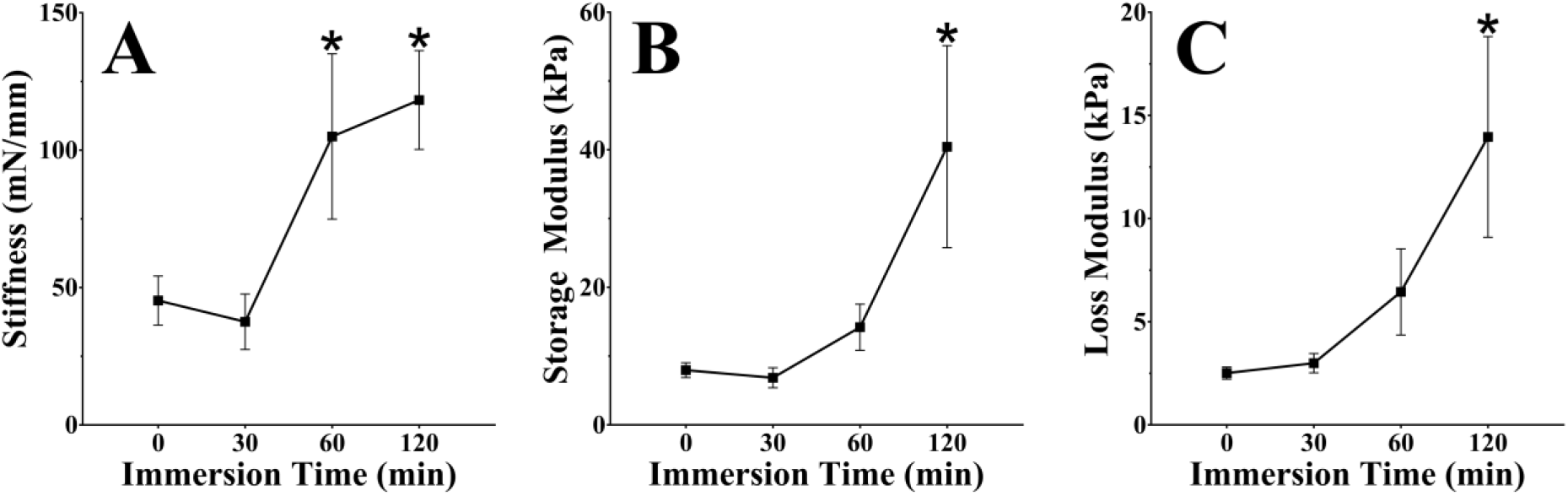
Effect of Ethanol Compaction on Scaffold Mechanical Properties. Graphs depicting **A**) stiffness, **B**) storage modulus and **C**) loss modulus of ABNP scaffolds tested after various immersion times in buffered 28% ethanol. * Indicates a statistical difference (p≤0.05) compared to 0 minutes.

### Dynamic Mechanical Analysis of Crosslinked and Compacted ABNP

To determine the combined effects of compaction and crosslinking on the dynamic mechanical properties of the ABNP, DMA was performed after various crosslinking treatments: conventional crosslinking (MES Only), ethanol-enhanced crosslinking (EtOH/MES), or ethanol-mediated compaction with enhanced crosslinking (Comp+EtOH/MES, Figure 4A). At 0.01 Hz and 0.1Hz, no significant differences in compressive stiffness were observed among experimental groups, although bNP and crosslinked groups were marginally stiffer than the ABNP (Figure 4B). At higher frequencies, these differences were more apparent; at 1 Hz, bNP and Comp+EtOH/MES scaffold stiffnesses were respectively 118.5±36.5 mN/mm and 149.1±43.3 mN/mm, which were significantly higher than the ABNP stiffness of 34.0±14.8 mN/mm (p<0.0287 and p<0.0020, resp.). At 10 Hz, bNP, MES Only, EtOH/MES, and Comp+EtOH/MES scaffolds demonstrated stiffnesses of 140.1±47.3 mN/mm, 124.1±46.6 mN/mm, 115.9±12.5 mN/mm, and 174.3±49.6 mN/mm, which were all significantly higher than the ABNP stiffness of 40.7±15.2 mN/mm (p<0.0104, p<0.0307, p<0.0409, and p<0.0004 resp.).

**Figure 4.**
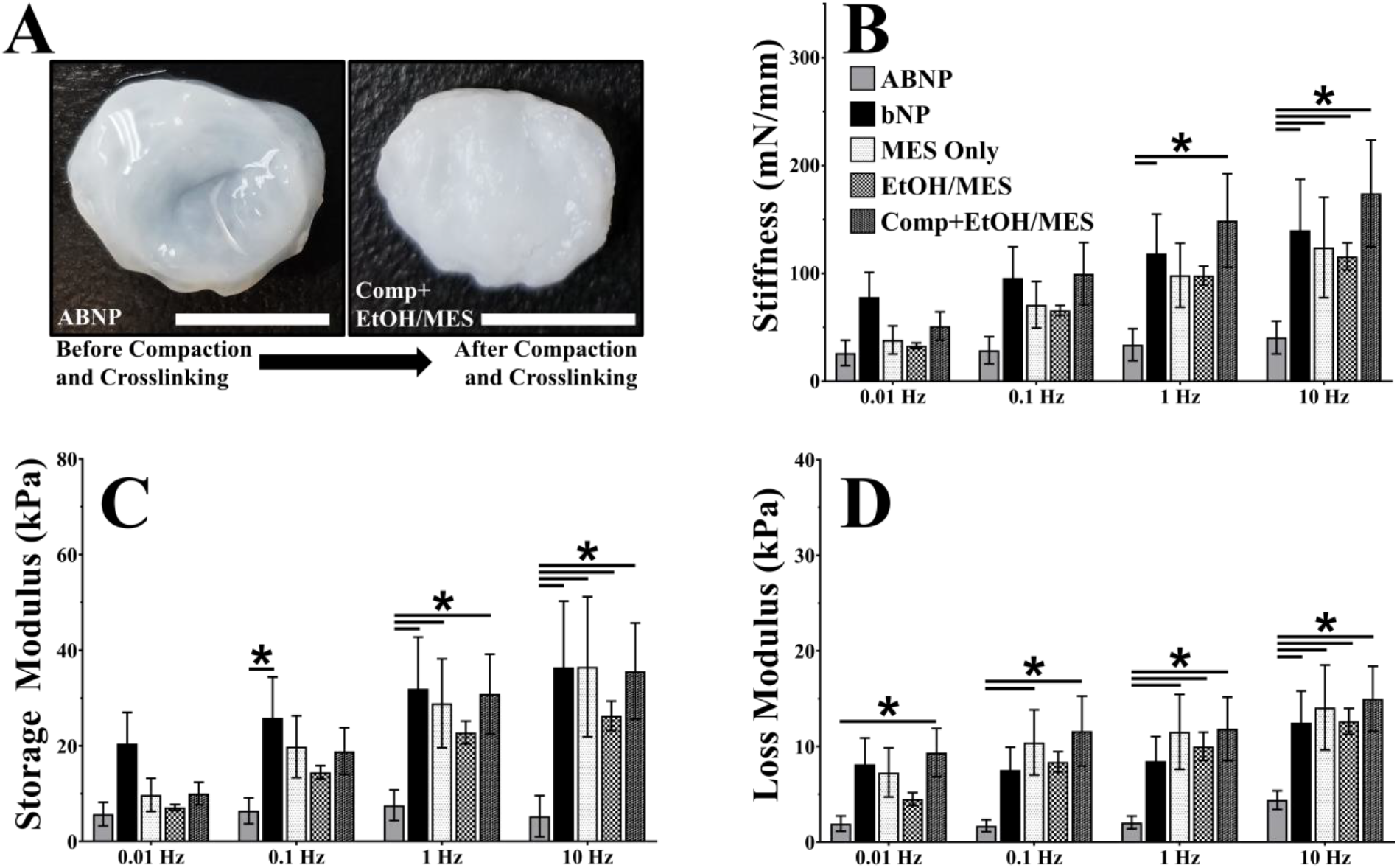
Scaffold Compaction and Crosslinking. **A**) Representative macroscopic images depicting a scaffold (left) prior to and (right) following compaction and crosslinking (bar = 1 cm). Graphs depicting **B**) stiffness, **C**) storage modulus and **D**) loss modulus of scaffold study groups tested at various frequencies. Lines connecting study groups indicate statistical differences (p≤0.05).

The trends in the storage moduli (Figure 4C) showed parallels to the compressive stiffness. At 0.01 Hz and 0.1 Hz, bNP and crosslinked groups had higher storage moduli than the ABNP. At 1 Hz, the ABNP storage modulus was 7.6±3.2 kPa. The bNP, MES Only, and Comp+EtOH/MES groups were significantly higher with respective storage moduli of 32.0±10.8 kPa, (p<0.0133), 28.9±9.3 kPa (p<0.0299), and 30.9±8.3 kPa (p<0.0131). Although falling short of significance, EtOH/MES storage modulus increased to 22.8±2.4 kPa (p<0.1019). At 10 Hz, ABNP storage modulus was 5.3±4.3 kPa, while bNP, MES Only, EtOH/MES, and Comp+EtOH/MES moduli were significantly higher at 36.5±13.8 kPa (p<0.0017), 36.5±14.7 kPa (p<0.0017), 26.3±3.1 kPa (p<0.0251), and 35.7±10.1 kPa (p<0.0.0014), respectively.

The loss modulus (Figure 4D) for ABNP at 0.01 Hz was 1.9±0.8 kPa, while bNP, MES Only, and EtOH/MES were marginally higher at 8.1±2.8 kPa, 7.3±2.6 kPa, and 4.5±0.7 kPa and Comp+EtOH/MES was significantly higher at 9.4±2.5 kPa (p<0.0316). At 0.1 Hz, loss modulus for ABNP was 1.7±0.6 kPa. bNP and EtOH/MES were higher at 7.5±2.4 kPa and 8.4±1.1 kPa, and MES Only and Comp+EtOH/MES were significantly increased at 10.4±3.4 kPa (p<0.0165) and 11.6±3.7 kPa (p<0.0045). The loss modulus at 1 Hz for ABNP and bNP were 2.0±0.7 kPa and 8.5±2.5 kPa, while MES Only, EtOH/MES, and Comp+EtOH/MES were significantly higher than the ABNP modulus at 11.5±3.9 kPa (p<0.0093), 10.0±1.5 kPa (p<0.0214), and 11.8±3.3 kPa (p<0.0049). At 10 Hz, the loss modulus for all groups compared to the ABNP was significantly higher, with ABNP, bNP, MES Only, EtOH/MES, and Comp+EtOH/MES groups having respective loss moduli of 4.4±1.0 kPa, 12.5±3.3 kPa (p<0.0256), 14.1±4.5 kPa (p<0.0080), 12.6±1.3 kPa (p<0.0174), and 15.0±3.4 kPa (p<0.0025).

### Compacted and Crosslinked ABNP Resistance to Accelerated Enzymatic Degradation

Exposing scaffolds to rhADAMTS-5 induced a mass loss of 30.5±2.4% in bNP and 28.8±7.8% in ABNP (Figure 5A). This was higher than the 6.1±3.0% loss in EtOH/MES group and significantly higher than the MES Only or Comp+EtOH/MES groups which respectively lost 14.3±3.6% (p<0.0158) and 3.2±3.2% (p<0.0076). Similarly, exposure to rhMMP-13 induced mass loss of 27.8±5.0% in bNP and 27.8±4.0% in ABNP, which was significantly higher than all crosslinked groups, which lost 10.9±6.1% (p<0.0437) for MES Only, 1.7±1.7% (p<0.0033) for EtOH/MES, and none detected (ND, p<0.0022) for Comp+EtOH/MES (Figure 6A).

**Figure 5.**
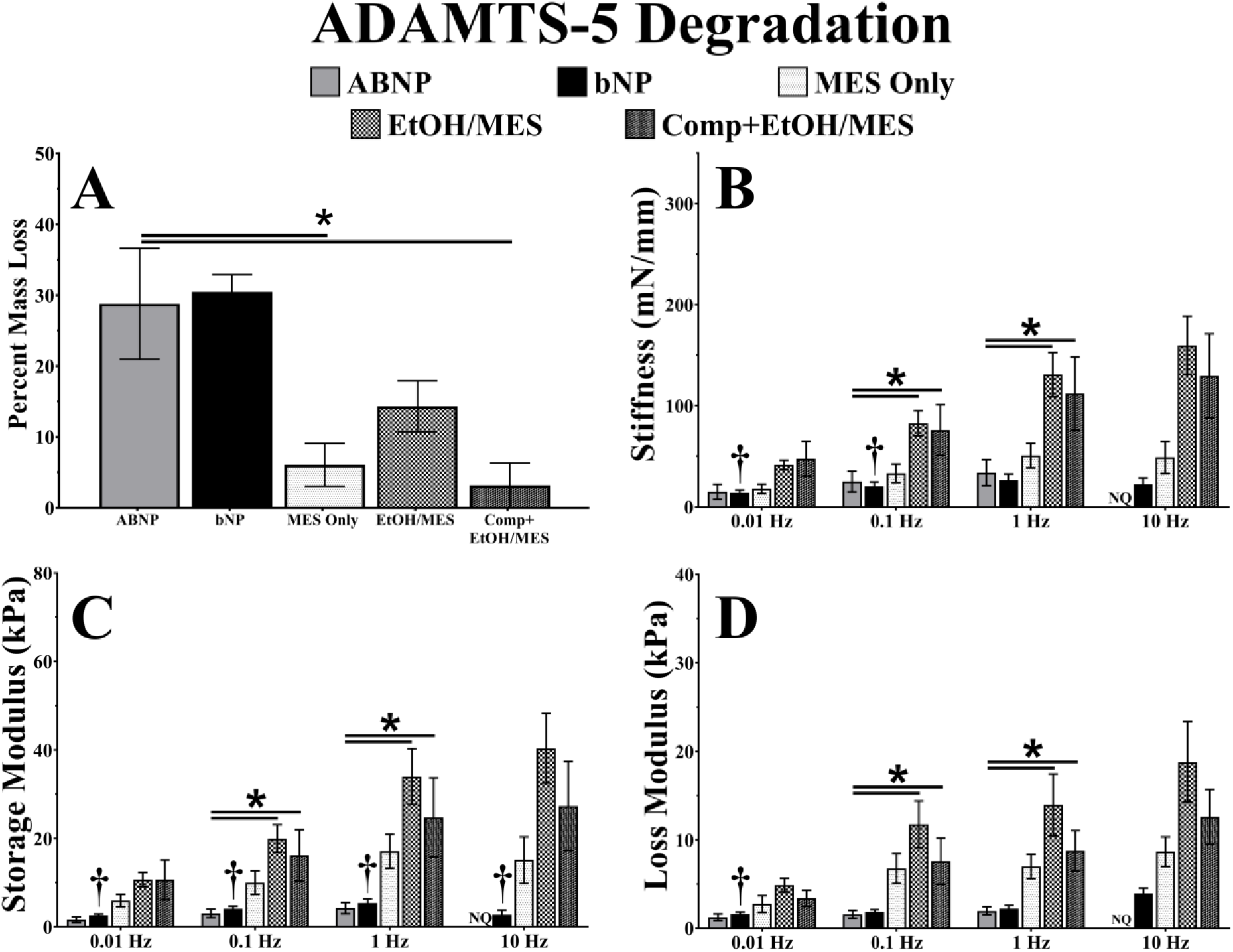
Scaffold Resistance to ADAMTS-5 Degradation. Graphs depicting **A**) mass loss and resultant **B**) stiffness, **C**) storage modulus, and **D**) loss modulus of scaffold study groups tested at various frequencies following incubation in recombinant MMP-13. Lines connecting study groups indicate statistical differences (p≤0.05), and † indicates statistical difference (p≤0.05) from pre-digested values. (NQ: not quantifiable).

**Figure 6.**
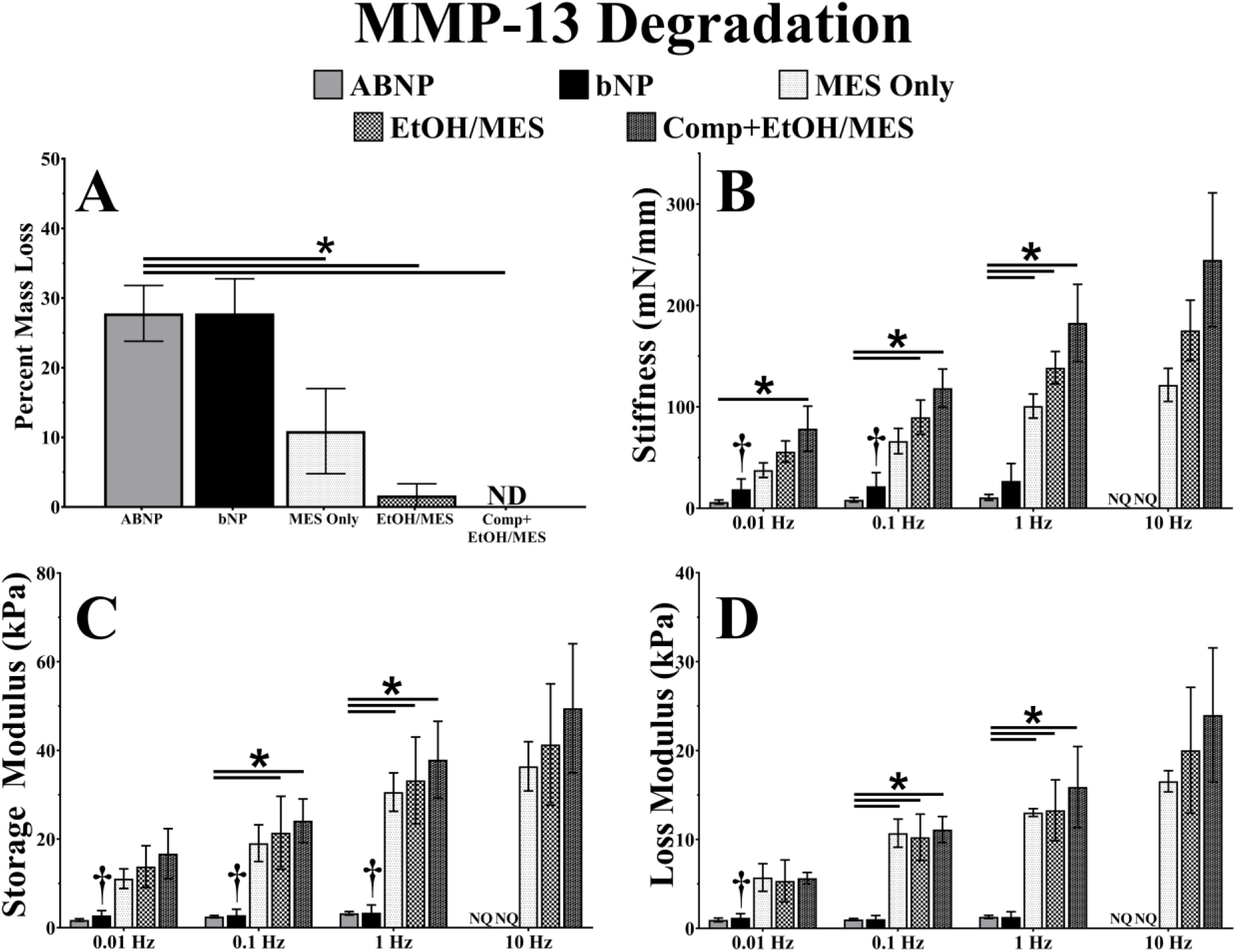
Scaffold Resistance to MMP-13 Degradation. Graphs depicting **A**) mass loss and resultant **B**) stiffness, **C**) storage modulus, and **D**) loss modulus of scaffold study groups tested at various frequencies following incubation in recombinant MMP-13. Lines connecting study groups indicate statistical differences (p≤0.05), and † indicates statistical difference (p≤0.05) from pre-digested values. (ND: none detected; NQ: not quantifiable)

To determine the ability of compaction and crosslinking to impart enzyme resistance, scaffolds were exposed to rhADAMTS-5 or rhMMP-13, followed by DMA. EtOH/MES scaffolds and Comp+EtOH/MES scaffolds were the only groups to maintain significantly higher compressive stiffness than the ABNP after ADAMTS-5 exposure at most frequencies (Figure 5B). On the other hand, rhADAMTS-5 had significant ramifications for the bNP group, whose stiffness significantly decreased from 78.1±23.0 mN/mm to 14.0±2.7 mN/mm at 0.01 Hz (p<0.0118) and 95.8±28.9 mN/mm to 20.5±4.1 mN/mm at 0.1 Hz (p<0.0448). Although not significant, decreases in stiffness were observed for the ABNP as well, and, at 10 Hz, the scaffold recoil was inadequate to quantify (NQ – not quantifiable) its DMA properties. Scaffolds crosslinked in MES Only tended to decrease in stiffness at each frequency following exposure to rhADAMTS-5. Exposure to rhMMP-13 similarly had a detrimental effect on bNP and ANBP compressive stiffness, which respectively decreased to 18.8±10.2mN/mm (p<0.0213) and 6.2±2.1 mN/mm at 0.01 Hz, 21.8±13.4mN/mm (p<0.0493) and 8.3±2.2 mN/mm at 0.1 Hz, and 27.0±17.2 mN/mm and 10.8±2.9 mN/mm at 1 Hz, and which were NQ at 10 Hz due to inadequate recoil (Figure 6B). In contrast, Comp+EtOH/MES scaffolds were significantly higher than the ABNP at all frequencies, EtOH/MES scaffolds and Comp+EtOH/MES scaffolds maintained their storage moduli at all frequencies after exposure to either rhADAMTS-5 (Figure 5C) or rhMMP-13 (Figure 6C). Furthermore, Comp+EtOH/MES storage modulus was significantly higher than that of the ABNP at more frequencies than any other test group. bNP storage modulus suffered significant decreases after digestion with rhADAMTS-5, reducing to 2.6±0.3 kPa at 0.01 Hz (p<0.0009), 4.2±0.6 at 0.1 Hz (p<0.0048), 5.4±0.9 kPa at 1 Hz (p<0.0138), and 2.8±1.0 kPa at 10 Hz (p<0.0205). rhMMP-13 had a similar effect on bNP, with a significant reduction to 2.8±1.1 kPa at 0.01 Hz (p<0.0009), 2.8±1.3 kPa at 0.1 Hz (p<0.0029), and 3.4±1.7 kPa at 1 Hz (p<0.0084), while it was NQ at 10 Hz. ABNP scaffolds, although not significantly decreased compared to their initial state, were markedly lower than crosslinked scaffolds at all frequencies for both enzymatic treatments. Trends in the loss moduli were comparable to the storage moduli, with bNP loss modulus tending to decrease at all frequencies after rhADAMTS-5 (Figure 5D) or rhMMP-13 (Figure 6D) treatment, significantly so at 0.01 Hz to 1.6±0.3 kPa (p<0.0184) and 1.2±0.5 kPa (p<0.0126), respectively. ABNP and MES Only moduli tended to decrease, and all crosslinked groups showed significantly higher loss modulus than the ABNP at 0.1 Hz and 1 Hz.

### Cytocompatibility of Crosslinked and Compacted ABNP

The Alamar Blue assay indicated that cellular activity increased for all crosslinked groups and was significantly higher than the ABNP at day 3, with the EtOH/MES increasing to 131.2±24.2% (p<0.0155), MES Only increasing to 115.1±3.8% (p<0.0003), and Comp+EtOH/MES increasing to 113.1±2.6% (p<0.0239) (Figure 7A). In contrast, the ABNP control group decreased to 88.0±5.7% at day 3. Cellular activity of all scaffolds declined at day 7, with ABNP decreasing to 78.0±2.8%, MES Only decreasing to 91.2±1.2%, EtOH/MES decreasing to 89.4±4.2%, and Comp+EtOH/MES significantly decreasing to 72.2±4.0%. Despite this decline in activity, no significant difference in cellular metabolism was found between the ABNP and any other group, and Live/Dead fluorescence (Figure 7B-G) showed that scaffolds retained large populations of viable cells with few dead cells at 10 days post-seeding.

**Figure 7.**
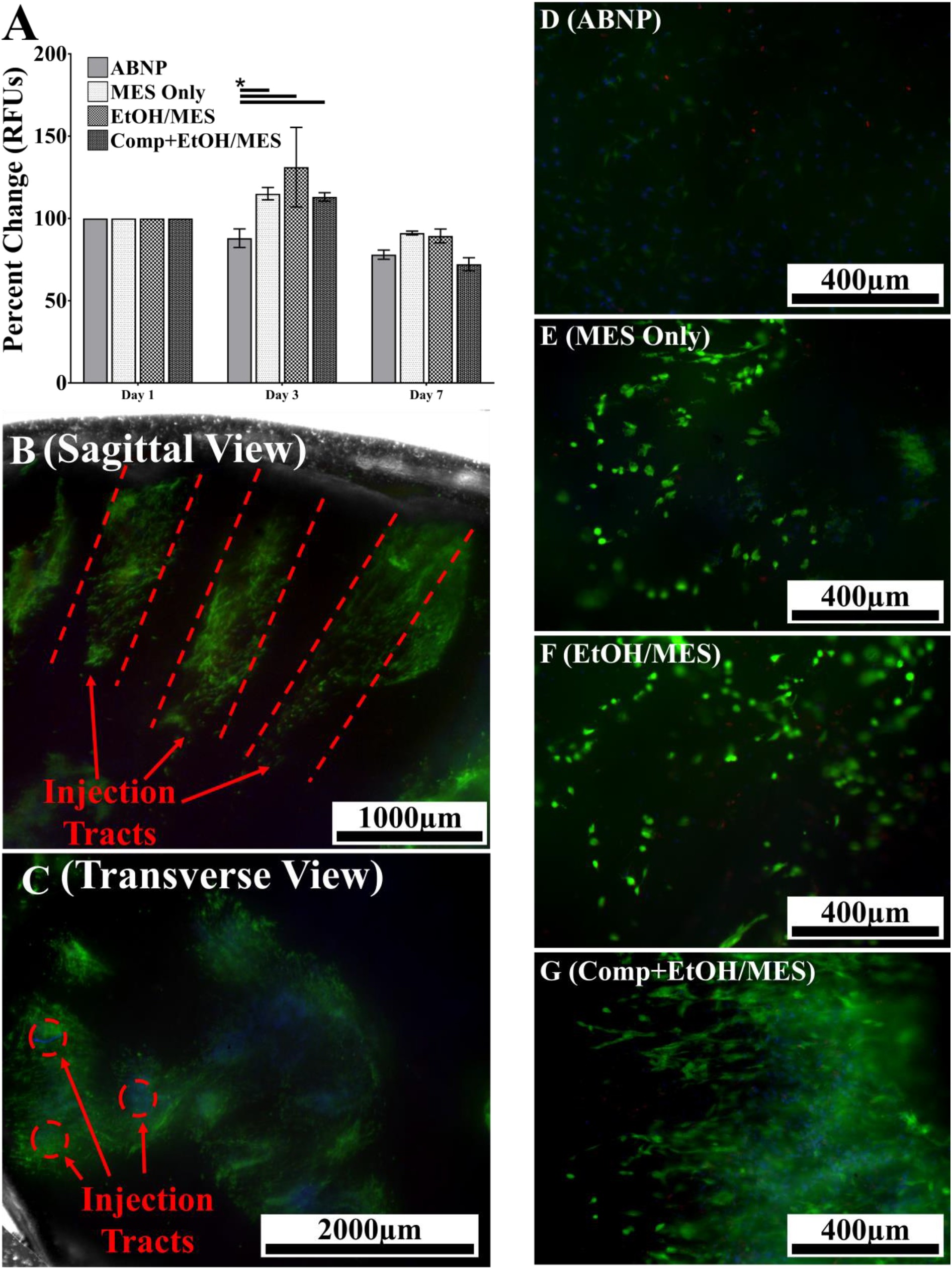
Assessment of Scaffold Cytotoxicity. **A**) Graphical representation of Alamar blue metabolic assay. Lines connecting study groups indicate statistical differences (p≤0.05). Representative low magnification LIVE/DEAD images of a scaffold in the **B**) sagittal and **C**) transverse planes illustrating the presence of viable stem cells (green) in the injection path (red dotted lines). Representative LIVE/DEAD images of **D**) ABNP, **E**) MES only, **F**) EtOH/MES and **G**) Comp+EtOH/MES illustrating overall cell viability (green = live cells, red = dead cells, blue = stem cell nuclei).

## Discussion

Herein, we evaluated the effects of ethanol-enhanced crosslinking on ABNP scaffolds. We demonstrated that MES buffer formulated from 28% EtOH induced compaction of our scaffolds, which had pronounced effects on the DMA properties of the ABNP, even in the absence of crosslinker. This finding was incorporated into a crosslinking methodology, which we then used to evaluate the effects of crosslinking, EtOH co-solvency, and EtOH-induced compaction. Supplementation with EtOH enhanced the mechanical properties of scaffolds, fortified them against enzymatic degradation, and demonstrated comparable cytocompatibility to the ABNP.

We have previously demonstrated that bovine caudal NPs can be effectively decellularized to produce human-mimetic NP scaffolds with similar ECM composition, mechanical properties, and cytocompatibility; however, the mechanical testing values were lower than those reported for human NP.^38^ This was in agreement with our bovine explant FSU model, for which the ABNP was not able to restore the viscoelastic creep parameters following repair.^39^ These mechanical deficiencies were attributed to decellularization-induced changes that increased ECM porosity and permeability due to free-swelling, GAG leaching, and tissue damage, which has been observed in other decellularization methodologies for NP tissue.^33,36,37^

To counteract the scaffold changes resulting from decellularization, we employed ethanol as a co-solvent for crosslinking. The use of co-solvents to control hydrogel swell state has been well studied using synthetic hydrogels,^45,46^ and it has been widely observed as a side effect of many fixatives used to preserves tissues for imaging.^47^ Using 28% ethanol buffered with MES, we reduced the scaffold volume to ∼70% of its original size (Figure 2). To our knowledge, this is the first attempt to induce compaction of a decellularized scaffold mediated by ethanol; however, Hiroki et al demonstrated that poly(acryloyl-l-proline methyl ester) hydrogels have a swelling capacity that can be controlled by alcohol concentration, inducing volumetric changes ranging from 0.5-14x change in gel volume in response to the alcohol fraction. Interestingly, Hiroki et al also found their constructs reached an equilibrium based on alcohol concentration, and that gels gradually decreased in size from 0-20% alcohol and reached their lowest swell state (50% of initial volume) from 20-43% alcohol.^45^ This aligns with our findings that 30% ethanol reduces the volume of our decellularized scaffolds to ∼70%.

In addition to the volumetric compaction, increases in the dynamic mechanical properties were observed in our acellular constructs. The DMA parameters increased 2-6x in response to ethanol compaction alone (Figure 3), which was comparable in magnitude to the bNP DMA parameters used for comparison later in the study. One explanation for this increase is the change in porosity with compaction. Perie et al demonstrated that increasing the NP porosity through free swelling or enzymatic degradation resulted in significant changes to the compressive modulus and permeability,^28^ which are related through the biphasic model described by Holmes and Mow.^48^

Interestingly, significant changes in the DMA properties were seen no earlier than 60 min and as late as 120 min following immersion in 28% EtOH, despite observing that the volumetric equilibrium was reached after 45 min. We attributed this to the highly porous nature of the ABNP and theorize that the initial changes in volume were likely sudden, but made insignificant contributions to reducing the hydraulic permeability; however, as the scaffold approached volumetric equilibrium, intermolecular resistance could have slowed compaction. Furthermore, at this highly compacted state, changes in ECM density may have had a pronounced effect on the hydraulic permeability. This could explain the preliminary responsiveness of the ABNP to volumetric changes without proportional mechanical changes, as well as the terminal increases in DMA properties with only minor changes in compaction. Additional contributions to the mechanical properties could have been made by ethanol, a coagulant fixative used in a variety of preservation protocols for microscopic imaging.^49^ Ethanol fixes tissue by altering the solubility of dissolved proteins, causing them to precipitate and agglomerate; however, this is an unlikely contributor since such fixation requires >50% ethanol.^50^

After evaluating compaction in isolation, we incorporated it into a crosslinking treatment for the ABNP scaffolds. Compaction and crosslinking significantly increased the DMA properties of the ABNP, was comparable to the bNP control, and tended to be marginally higher than other crosslinked groups (Figure 4). We also observed that the mechanical measures of our compacted and crosslinked samples bore similarities to human values reported by Freeman et al, who showed the human NP to possess respective storage and loss moduli of 25–125 kPa and 10–47 kPa, in comparison to 10-36 kPa and 9-15 kPa seen in our samples.^51^ Additionally, the improved DMA properties resulting from compaction and crosslinking did not have detrimental side-effects on scaffold cytotoxicity, as evidenced by comparable cellular activity throughout and viable cells up to 10 days post-seeding (Figure 7).

The biologic origin of our acellular scaffold makes it susceptible to proteases known to be active in IDD;^44^ as an orthopaedic implant, the ABNP must be able to tolerate this catabolic environment while maintaining its mechanical functionality *in vivo*. Thus, compacted and crosslinked samples underwent DMA before and after exposure to an accelerated-degradation model mediated by recombinant human ADAMTS-5 (Figure 5) or recombinant human MMP-13 (Figure 6). We observed that EtOH-supplemented groups tended to resist degradation, which was demonstrated by maintained DMA parameters following degradation and lower mass loss. This is in contrast to the bNP and ABNP samples, whose stiffness, storage modulus, and loss modulus were reduced by 2-12 fold, and whose mass loss was significantly higher than that of any crosslinked group. Our findings were mirrored those of Perie et al, who showed that swollen bovine NP subjected to digestion with proteolytic enzymes exhibited an approximate 10-fold decrease in compressive modulus.^28^ Herein, we have demonstrated the ability to protect the native bovine NP ECM present in our scaffolds via ethanol-supplemented crosslinking.

### Limitations

The research herein was performed with certain limitation. Introducing compaction into our crosslinking treatment likely induced changes in the biphasic properties of our constructs, which we quantified via dynamic mechanical analyses. Confined compression could provide a more direct evaluation of these changes and has been well described in the literature for NP tissue.^52–55^ Another limitation is our implementation of the enzymes used for degradation, which was based on the concentrations thought to occur in a degenerative environment rather than their activity levels.^44^ With batch-to-batch variations in enzyme activities, it is possible that our simulated degradation could have over-or under-estimated that seen in vivo; however, patient-to-patient variability could result in similar discrepancies, and, since all samples received uniform treatment, the relative changes remain valid. An explanation and limitation for the decreasing cell metabolism observed in all samples could be the high passage number of our initial seeding population (passage 6), which has been shown to induce detrimental alterations to stem cell differentiation and senescence.^56,57^ However, by seeding scaffolds from the same pool of cells, normalizing metabolic activity to day 1 values, and using the starting material (ABNP) as the control, this limitation was minimized.

## Conclusions

Herein, we demonstrated that incorporating ethanol as a crosslinking co-solvent for our ABNP implant enhanced its dynamic mechanical properties and resistance to enzymatic degradation without altering the scaffold’s innate cytocompatibility. Furthermore, we attributed ethanol’s effects to its induction of volumetric compaction of the scaffold and improved crosslinking efficiency. Future mechanical testing must be performed to further verify these effects, and whether they have long-term ramifications on seeded stem cells. Future work also includes developing strategies to incorporate additional ‘aggrecan-mimetic’ analogs to compensate for those lost during decellularization to further enhance the mimetic nature of this scaffold.

## Acknowledgements

This research was supported by the National Institute of General Medical Sciences of the National Institutes of Health (award number: 5P20GM103444) and the Technology Maturation Fund awarded by the Clemson University Research Foundation.

